# Optimization of adhesion for high throughput cryo-electron tomography of vitreous sections

**DOI:** 10.1101/2025.10.09.681450

**Authors:** Fatima Taiki, Helmut Gnaegi, Mikhail Eltsov, Amélie Leforestier

## Abstract

Cellular cryo electron tomography explores tissue and cells in their unstained flash-frozen native state, revealing *in situ* the structure of macromolecules together with their local environment and interactions with partners, also known as molecular sociology. To obtain thin samples, cryo-FIB milling is nowadays the most popular method, with impressive successes. The alternative, cryo-ultramicrotomy, is often overlooked on account of poorly reproducible attachment of cryo-sections to their support, resulting in extremely low throughput. We optimized the workflow, focusing on section adhesion and their support grids. We thus increased vitreous sections’ cryo electron tomography throughput to equal that of thin film, with typically several tens to hundreds of cryo-tomograms per sample. This open the way to new advances in cellular cryo electron tomography, as the method is devoid of beam damage, can provide large surfaces and serial sections of any type of sample from cells to tissues. In addition, section thickness can be tuned down to 30-50 nm, which may be an advantage for imaging small molecular complexes, such as DNA and nucleosomes.

## INTRODUCTION

Cellular cryo electron tomography is experiencing continuous spectacular developments, since its opening of the field of molecular sociology (Robinson et al., 2007), revealing with increasing resolution and complexity the structures and interactions of macromolecules at work inside the cell (Nogales and Mahamid, 2024). These achievements have been possible thanks to new generations of high-end stable and automated electron microscopes and direct electron detectors, and the development of cryo-FIB milling to achieve sample thinning to electron transparency,i.e. less than a few hundred nanometers, ideally below 200 nm (Villa et al., 2013) (Rigort and Plitzko, 2015). The historic alternative to cryo-FIB milling is cryo-sectioning – also known as CEMOVIS (cryo electron microscopy of vitreous sections), which cuts samples into thin slices with a diamond knife. CEMOVIS offers a number of advantages. (i) It is able to provide ultra-thin samples, with sections as thin as 20 nm (Ng et al., 2020), more routinely in the 40-80 nm range, whereas cryo-FIB lamellae are usually thicker than 100 nm, routinely in the 150-200 nm range. This may be an advantage when high resolution is needed or to image small complexes and molecules, such as DNA (Fatmaoui et al, 2025). (ii) Cryo-sectioning provides access to large, entire volumetric domains of a specimen. Not only the typical section area is about 20 times larger than a cryo-FIB lamella, but cryo-sectioning generates serial sections of a sample, whereas FIB-milling destroys most of it, keeping only one lamella – or a few sparse ones (Schiøtz et al., 2024). In addition, cryo-sectioning can be applied equivalently to any type of sample, cell suspensions or tissues, after high pressure freezing. If cryo-FIB milling of cells that can be frozen on EM-grids is now a high throughput technique, it is much more challenging for tissues, for which complex lift out methodologies are required (Parmenter and Nizamudeen, 2021). Lastly, cryo-FIB milling is itself not artefact-free, and is in particular associated to ion beam damage that can extend to a depth of 60 nm from the surface, limiting the recovery of information therein (Lucas and Grigorieff, 2023), which is not the case with cryo-sectioning (Elferich et al, 2025). A few studies have indeed highlighted CEMOVIS potential for high resolution imaging, with structural information retrieved down to 2.9 Å (Moriscot et al., 2023) or 7.9 Å (Sader et al., 2009), or 3D structure reconstruction at sub-nanometer resolution, down to 3.5 Å and even 2.98 Å on ribosomes subunits (Elferich et al, 2025)(Al Amoudi et al., 2005), very similar to what can be obtained from FIB-milled samples (Hoffmann et al., 2022). However, CEMOVIS is nowadays overlooked, on account of compression artefacts induced by the cutting process (Alamoudi et al., 2005), but also because of the extremely low throughput of the method, especially for the recording of tilt series for cryo-tomographic reconstructions. The poor yield in recording stable tilt series on cryo-sections is primarily due to poorly and irreproducible adhesion of cryo-sections to their support grid, but also to the difficulty in assessing perfect adhesion *a priori*, before the recording of the tilt series themselves. In addition, for cryo-ET imaging in high end microscopes, grids must be clipped, which can result in breaks and loss of sample areas. This is rarely a problem for plunge-frozen samples whose support film is reinforced by its covering with the layer of frozen buffer solution, but becomes an issue when working with cryo-sections simply deposited on a thin support film.

In this study, propose an optimized workflow leading to high throughput cryo-ET recording. We introduce an extra step to improve adhesion, using the electrostatic press developed for section attachment by Pierson et al (2011), away from the knife edge. We also propose a global real-time monitoring test of adhesion upon section collection on grids, and a local adhesion characterization in low magnification mode before image/tilt series recording. Lastly, we test a variety of support grids, and determine optimal supports for either 2-dimensional (2D) or 3-dimensional tomographic imaging.

## MATERIAL AND METHODS

### Sample vitrification

Stage 13-15 Drosophila embryos were dechorionated in 50% (v:v) bleach, rinsed in PBS and transferred into a Leica type A gold-plated copper carrier. The carrier was then filled with 25% (w:w) dextran (40 kDa, Sigma-Aldrich) and 0.25% of NP-40 (Sigma-Aldrich) in PBS and covered with the flat side of a Leica type B aluminum carrier. The sandwich was transferred into a HPM010 freezing machine, and vitrified with liquid nitrogen at 2045 bars.

### Cryo-sectioning

High-pressure frozen samples were transferred into the chamber of a FC6/UC6 cryo-microtome (Leica Microsystems) installed in a controlled environment room, with the relative humidity set to 20-25 % and the temperature to 23°C. The cryochamber of the microtome was cooled down to –140°C. The aluminum cover was removed and the gold-plated copper carrier filled with the sample mounted inside a flat sample holder. A diamond trim 45° (Diatome) was used to remove the metal edges of the carrier, and a ∼ 50 μm x 70 μm x 50 μm pyramid was trimmed within either the yolk or the central nervous system of the embryo. Final trimming steps were realized using a 20° trim knife (Diatome). Cryosections were cut from the pyramid at a nominal feed of 40, 50 or 75 nm with a cryo 25° diamond knife (Diatome) at a cutting speed of 0.4 mm/sec. A Leica EM Crion tunable antistatic device (static line) in discharge mode was use to optimize section generation and gliding along the knife. As recalled in Figure 1 (A, B), ribbons of serial sections were pulled and collected using a double micromanipulator (Leica), which lets one control the obtention of millimeter-long ribbons of sections (Pierson et al, 2011) (Studer et al., 2014). The sections were then collected onto support grids. Adhesion is obtained using the Leica EM Crion antistatic device positioned perpendicular to the grid, used in charging mode, which acts as an electrostatic press (Pierson, 2011).

**Figure 1.**
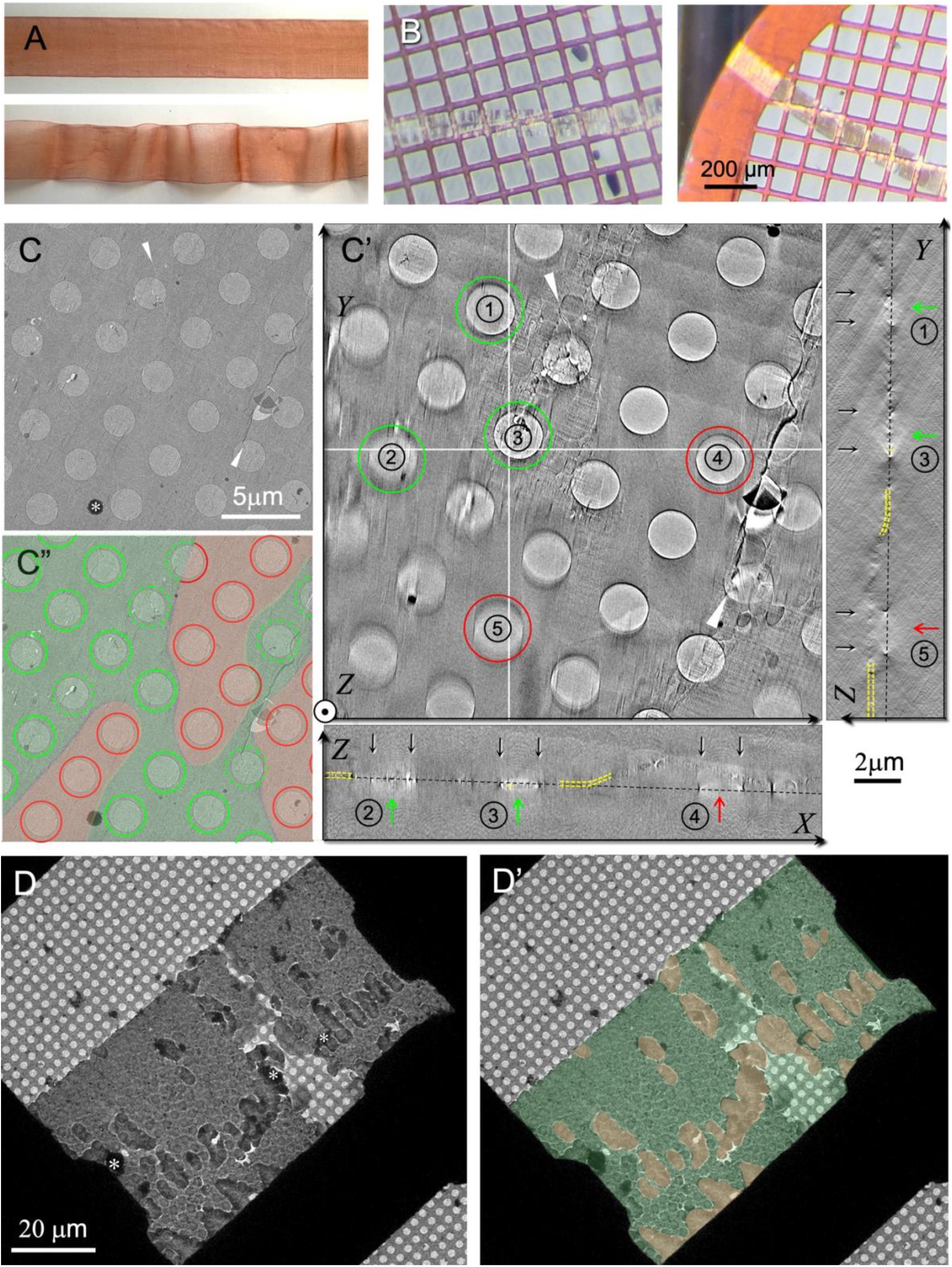
Adhesion tests. (A) Flat ribbon in close contact with its support (left); krinkled ribbon with sparse contacts (right). (B, B’) Adhesion revealed at the macroscopy level: the cryo-section ribbon turns, initially silvery turns gold-purple after charging with the electrostatic press (B’). Non-adhering ribbons remain silvery (B). (C) Low magnification imaging of a section. White arrowheads point to breaks in the section. White asterisks indicate ice contamination deposited on the section. (C’) Low magnification cryo-tomographic reconstruction of the region shown in (C). Virtual slices in XY, XZ and YZ are shown. Holes (1), (2) and (3) (circled in green) are located in adhesive regions in tight contact with the carbon film. The section is floating above holes (4) and (5) (circled in red) located within a non-adhesive area. In XZ and YZ slices the carbon film is indicated by the black dotted line. The profile of the cryosection is locally underlined by yellow dotted lines. Hole boundaries are pointed by the black arrows. (C”) Mapping of the area shown in (C) as determined by the tomographic reconstruction. Adhesive areas are coloured in green, non-adhesive ones in red. Holes circled in green can be used for high magnification tomographic acquisitions (the dotted line indicates that, despite located within an adhesive region, the area is not optimal (presence of breaks or partially adhesive border). (D, D’) Region of an atlas acquired at 5 mm defocus. A differential contrast is observed between adhesive (coloured in green in E’) and non-ashesive (in red) regions. Supports: (B, C-C”) Quantifoil R2/2; (D, D’) Quantifoil R2/1 reinforced with 30 nm carbon.

### Support grids

We tested different support films: Quantifoil R2/1, R2/2, R3.5/1 and S7/2, UltrAuFoil R2/2 and C-flat CF-2/1, all copper or gold (UltrAFoil) 200 mesh grids (Electron Microscopy Sciences). Some of them were covered with a ultrathin (2-3 nm) continuous carbon film either commercial (Quantifoil R7/2), or evaporated on freshly cleaved mica from a carbon thread (Leica) using an ACE600 evaporator (Leica Microsystem), floated on distilled water, and deposited onto the immersed grids (Passmore and Russo, 2016). Some were reinforced with a 30 nm-thick extra layer of carbon evaporated directly on the back side of the grids. The different supports are listed and sketched in Figure 3.

### Cryo electron microscopy & tomography

For 2D imaging grids were mounted in a Gatan 626 cryoholder and transferred in a JEM-2010F-CRYO (JEOL) electron microscope operated at 200kV, equipped with a 4K Ultrascan 1000 CCD camera (Gatan). Images were acquired in “search” mode and minimal irradiation (≤ 0.001 e^-^/Å^2^) or in “photo” mode at a pixel size of 0.354 nm/pixel, with an electron dose of ∼10-15 e^-^/Å^2^, and a defocus of 2 or 3 µm. For cryo-electron tomography, grids were mounted in Autogrid rings (Thermo Fischer Scientific) and transferred into a Titan Krios (Thermo Fischer Scientific), operated at 300 kV and equipped with a Volta phase plate (VPP), a GIF Quantum SE post-column energy (Gatan). Images and tilt series were recorded with a K2 Summit direct electron detector (Gatan). Atlases were recorded at low magnification (pixel size 129.7 nm) and 2.5 to 5 mm defocus. Ultra-low dose tilt series were recorded at ×1400 (pixel size 11.33 nm) magnification using Serial EM software (Mastronarde & Held, 2017). Final tilt series recording was performed at a nominal magnification of ×64000 (2.12 Å/pixel), following the dose-symmetric recording scheme (Hagen, Wan et al., 2017) starting from 0°, with an angular range between -60° and +60° and a 2° increment. They were collected with VPP in close to focus conditions (0.25 μm defocus), with an electron dose set to 2.5 e^-^/Å^2^ /image.

### Image denoising. Tomogram reconstruction and denoising

Some of the 2D images were denoised using wavelet filtration in imageJ (“A trous” plugin with k_1_ = 20, k_n>1_ = 0). Low magnification low dose tomograms were reconstructed with the etomo program of IMOD (Kremer et al., 1996). For reconstructions of low magnification-low dose tomograms, a SIRT-like filter, equivalent to 10 iterations, was applied. Denoising of high magnification tomograms was performed with the deep learning network available in the Warp (Tegunov & Cramer, 2019) software package. For training data, dose-fractionation frames of the tilt series were aligned using the IMOD Alignframes tool, resulting in three tilt series: the complete one consisting of tilt images containing all frames for each tilt angle (*complete*); a tilt series containing even frames for each tilt angle (*even*); and a tilt series containing odd frames for each tilt angle (*odd*). Tilt series alignment was performed on the *complete* series using the patch tracking of etomo, and used to align the *odd* and *even* series. Tomograms were reconstructed in etomo using weighted back projection with the standard ramp filter. The Noise2Map network of Warp was trained on the *even* and *odd* reconstructions, starting with the initial model noisenet3dmodel_256 using the following parameters: dont_augment, dont_flatten_spectrum, learningrate_start 1E-05, iterations 40000.

## RESULTS

### Assessment of global (macroscopic) and local (microscopic) adhesion

Ideally, collected ribbons should be flat and fully adhere to their support; in real life sections may present pleats and/or adhere only sparsely to their support (Figure 1A), or, in worst cases, not at all. Sections obtained with last generation diamond knives using a clean, free of ice crystal contamination, antistatic device glide smoothly on the knife’s surface, leading to globally flat sections. Pleats may form during section generation, but can at least partly be removed upon gentle pulling during ribbon generation. If pleated regions remain in the final sample, they are readily identified upon low magnification observation and can be disregarded. But the observation of a flat region does not mean that the section adheres to its support. As an example, such a typical low magnification view of a cryosection is shown in Figure 1C. A 2D image recorded with a typical 10-20 e^-^/Å^2^ on a non-adhesive region generally suffers from drift and unproper defocusing. A tilt series recorded on a non-adhesive region usually moves independently of the underlying support tracked by the automated tilt series acquisition procedure, and a tomogram cannot be properly reconstructed. It is therefore essential, for optimal throughput, to sort out adhesive and non-adhesive regions beforehand. To check adhesiveness, we established different tests:

i. The global (macroscopic) adhesion of a ribbon on its support grid can be assessed already in the cryo-ultramicrotome chamber, using the electrostatic press. Ultrathin sections typically present a silver reflection color, expected for 50-80 nm-thick sections. Close contact between sections and their support films upon charging results in a change of light reflection: non or poorly adhesive sections remain silver, whereas fully adhesive ribbons become shiny and purple-gold (Figure 1B). This phenomenon is due to the change of thickness of the reflecting object: adhering sections “melt” with their carbon support, leading to a single thicker reflecting object, which shifts the reflection color. Samples which do not show this transition have too few or no adhesive regions, and are not suitable for high throughput tomography. This test is to be used to select best ribbons for tomography. However, it does not mean that the entire surface is adhesive, and it should be completed with local analysis before tilt series acquisition.
ii. Low magnification (typically x1400) recording of a tilt series with ultra-low dose irradiation of the specimen let us reconstruct the 3D shape of the section and determine its adhesion profile on its support film (Figure 1C-C”). Adhesive regions of the section appear in close contact with the supporting film (Figure 1C’, C”, green), while non adhesive ones are floating above (Figure 1C’, C”, red). Adhesives and non-adhesive regions can thus be sorted out and high magnification acquisitions can be targeted to adhesive regions (green circles in Figure 1C’, C”). However, this procedure is time consuming as a low magnification tomogram must be recorded and reconstructed for each region of the sample.
iii. Atlas recording with an ultra-high defocus, typically between 2.5 to 5 mm leads to a different contrast between adhesive and non-adhesive regions (Figure 1D, D’), which could correspond to a difference in phase shift. In any case, adhesive and non-adhesive regions can thus be told apart at a glance. We tested the phenomenon and its reproducibility by acquiring tilts series, which showed a perfect matching: among 206 tilt series acquired using this method, 175 turned out suitable for high quality tomographic reconstruction (mean residual in etomo < 1.0 and further analysis).
iv. At the microscopic local level, perfect adhesion of a section to its support film can also be revealed upon focusing the electron beam on the sample: adhesive regions bubble but do not melt (Supplementary Figure S1). In contrast, the focused beam creates a hole that propagates rapidly in non-adhesive regions, as it does on unsupported vitreous films. As this test is destructive, it cannot be used directly on ROI when a continuous ultrathin support is used. It is also very local and does not provide a complete mapping of adhesive/non adhesive regions. We use it for either for rapid test of adhesive properties of different support materials (see below) or when only 2D images are needed.

### A two charging steps procedure for improved section attachment

We recall in Figure 2A, B the experimental procedure to generate and collect a ribbon of cryo-section using a double micromanipulator, which lets one control the generation and collection of millimeter-long ribbons (Studer et al., 2014). Note that ribbons typically longer than 1.5 mm tend to twist or bend (Supplementary Figure S2). To attach the ribbon to its support grid, electrostatic charging, with the use of an electrode (Leica EM Crion) positioned perpendicular to the EM grid supporting the ribbon is now preferred to mechanical pressing, as it provides highest efficiency without risk of damaging the sample (Pierson et al., 2010). To optimize the Crion charging efficiency and attachment of the sections, we introduced a two-steps charging procedure sketched in Figure 2C: upon collection onto the support grid, a first electrical field is applied by the electrode, leading to the apparition of small purple-gold areas, indicative of locally adhering regions; then the grid is pulled away from the knife edge and rotated so that it lies perpendicular to the electrode; a second electrical field is finally applied before the grid is transferred out. Under this second charging step, the small adherent purple-gold areas initially visible expand. The ribbon is flattened against its support film, and the melting process propagates. Depending on electrostatic conditions, the charging pulse might need to be repeated, to maximize the contact areas. The complete attachment is not possible when the sections are not perfectly flat initially, because of geometric constrains: once an area is attached, it becomes unable to move and may prevent flattening and attachment of neighboring regions. A comparison of ribbons obtained with this procedure or with a simple single step charging is shown in Supplementary Figure S3. Note that an optimal electrostatic press is required to obtain these results: the electrode should be checked beforehand and cleaned regularly using a fiber glass brush tool (Radiospare). The decrease of press efficiency (with the aging of the electrode for example), can lead to weaker adhesion, which is revealed at the macroscopic level by the absence of change of light reflection upon use of the press (Figure 1B, B’).

**Figure 2.**
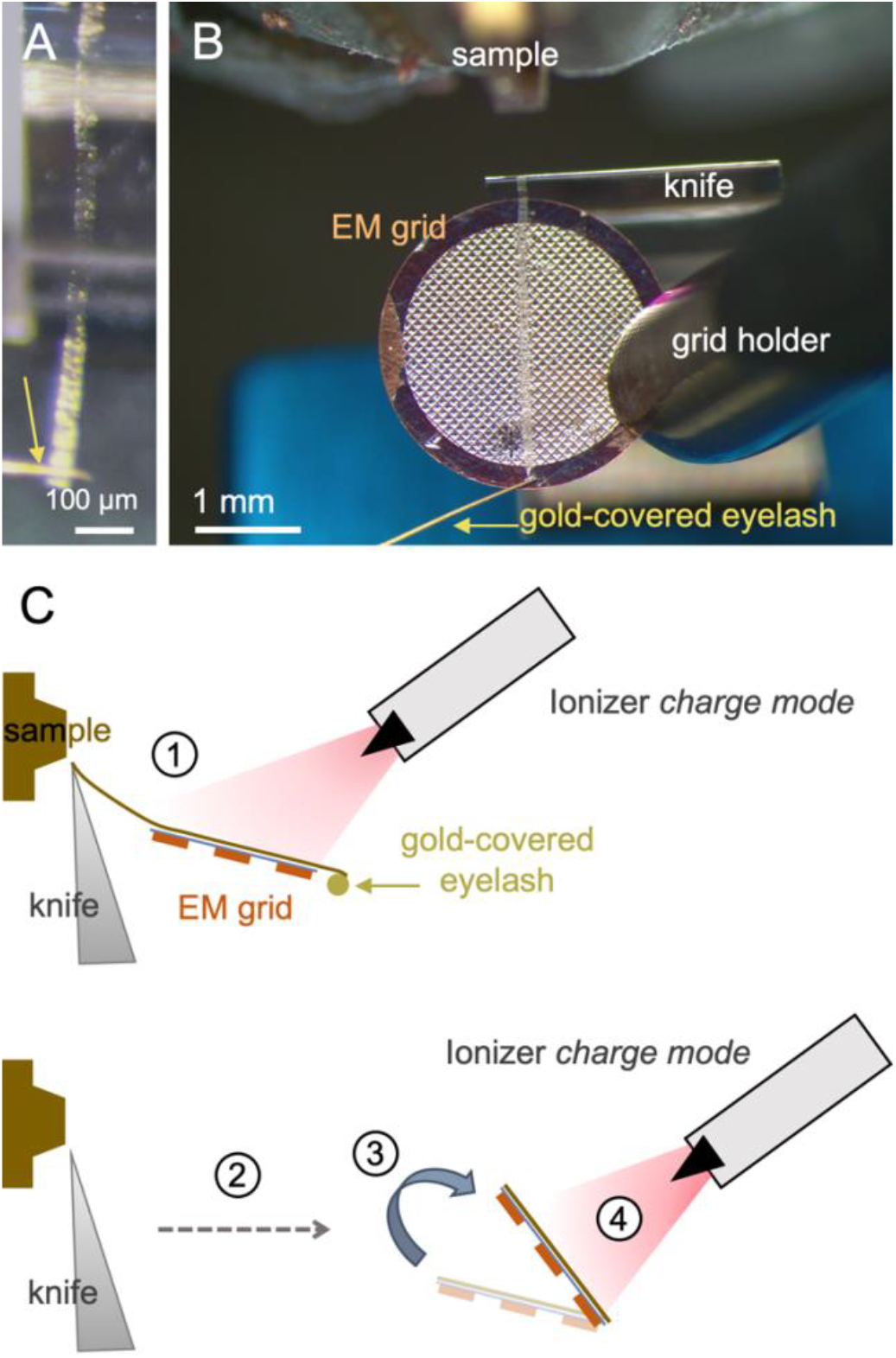
(A, B) Obtention and collection of ribbons of cryo-sections. (A) Ribbon of serial sections pulled with a gold-covered eyelash (golden arrow) controlled with a double micromanipulator in the cryochamber of the microtome. (B) Collection of the ribbon on its support grid. (C) Sketch of ribbon collection. A first “charge” electrical field is applied after contact grid the grid (1). Adhesion is optimized by displacement away from the knife edge (2) and reorientation of the EM grid perpendicular to the antistatic device (3) before applying a second “charge” ionization step (4).

### Test of different support films for optimization of section adhesion and resistance to clipping

In practice, different section supports have been used so far to collect cryo-sections, from thin continuous carbon films (Dubochet and Sartori Blanc, 2001), to carbon-covered formwar films (Hsieh et al., 2002) and lattices of holes in carbon films (Bharat et al., 2018) (Eltsov et al., 2018). They must fulfill several requirements which may sometimes be antagonistic: resistance to clipping, no or minimal added noise on recording areas, adhesivity, maximization of surface available for imaging. To our knowledge, there has not been a systematic investigation of any of these properties.

We considered different grid types, listed in Figure 3. We focused on carbon and gold films, principally different types of holey lattices:

**Figure 3.**
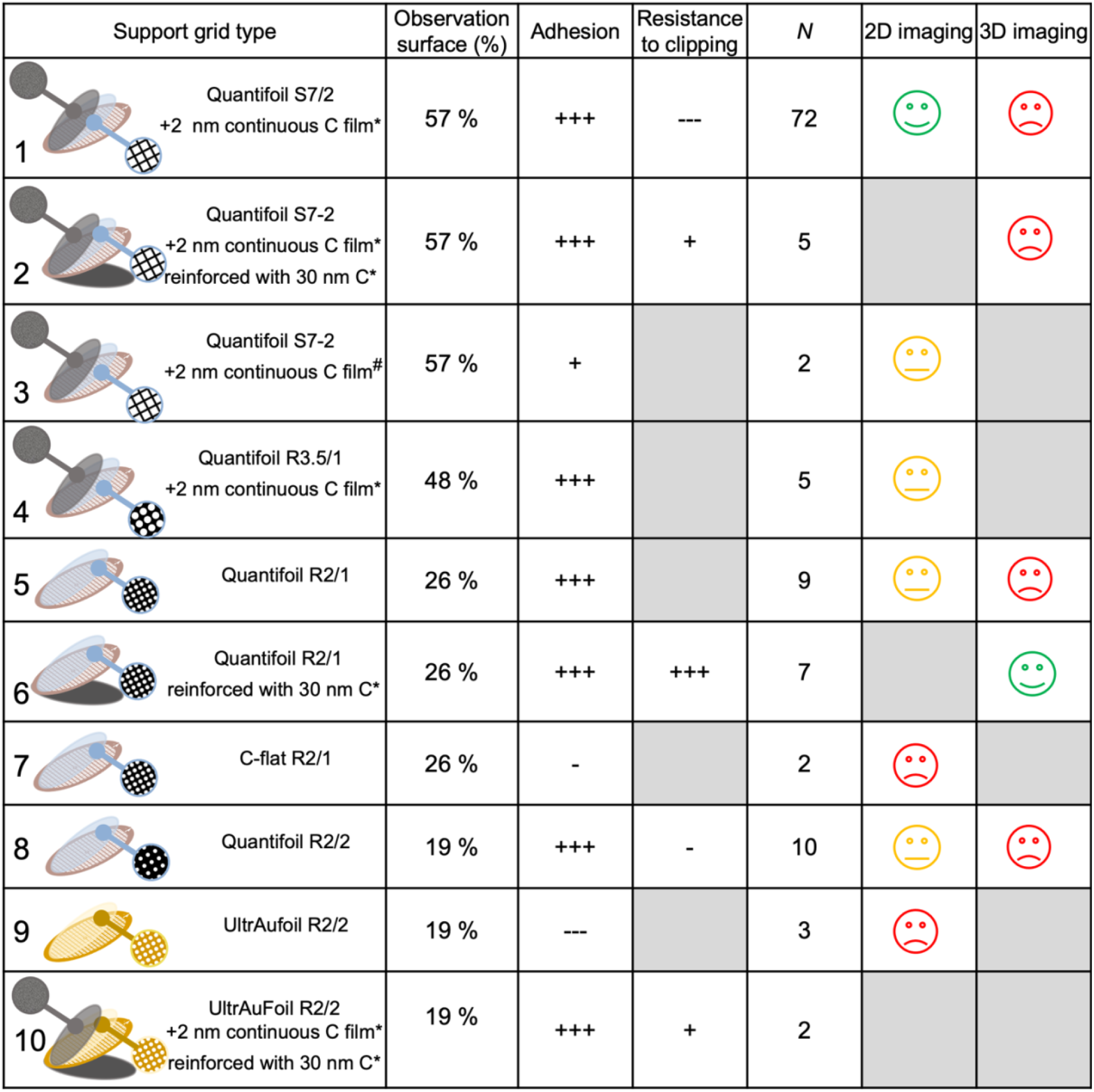
Characterization of different support grids for optimal cryo-EM imaging in 2D or 3D (tomography). *Additional carbon layers evaporated in our Leica ACE. # Additional carbon layer from furnisher (Quantifoil). Boxes appear in grey for conditions not tested. 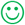 indicates the best performing supports, 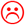 the worst ones to be avoided, and 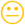 support films that can be used but are not optimal (slightly less adhesive, and/or lower available surface for imaging).

‐ Quantifoil round hole carbon arrays R2/1, R2/2, and R3.5/1
‐ Quantifoil square holes carbon arrays S7/2
‐ C-flat round hole carbon arrays R2/1
‐ UltrAuFoil round hole gold arrays R2/2

Too large areas of free-standing sections lead to increased instability under the beam. Large size hole arrays (S7/2 and R3.5/1) were therefore covered with a continuous ultrathin 2 nm carbon film, either homemade, evaporated in our Leica ACE evaporator (Figure 3^*^), or commercially available (Quantifoil S7/2, Figure 3^#^). We searched to optimize resistance to clipping by testing two different film thicknesses: commercial grids were thus used either directly - with their original 20 nm-thick film – or after evaporation of an extra 30 nm strengthening carbon layer, leading to a total thickness 50 nm (Figure 3, Supplementary Figure S4). For each condition, we tested from 2 to 72 sample grids (Figure 3, N).

Adhesion tests described above were performed on ribbons deposited on the different types of grids. The results are synthetized in Figure 3. Macroscopic adhesion revealed by change of light reflection upon contact between the grid and the ribbon was observed for samples collected on carbon surfaces from Quantifoil or homemade, but not for C-flat carbon surfaces and gold surfaces (UltrAuFoil), suggesting a poorer adhesion. Systematic microscopic adhesion tests were then realized either by beam focusing on sections regions covering the support film (Supplementary Figure S1), and/or high defocus low magnification mapping (Figure 1D, D’). In the first case, tests were followed by acquisition of 2D images; in the second, atlas recording was followed by acquisition of tilt series. Microscopic tests confirmed our macroscopic observations. 2D images acquired randomly on sections deposited on homemade carbon surfaces result in about 60-80 % of high-quality images - as characterized by the absence of drift. In good agreement, surface measurement on atlases recorded at high underfocus show that from about 50 to 80 % of a ribbon’s surface is adhesive on these films. In contrast, less than 10 % of high quality 2D images were obtained from samples deposited on C-flat carbon surfaces. We did not attempt to record tomograms on C-flat grids. Gold surfaces are the less adhesive ones, with nearly no adhering region found. No image or tilt series could be recorded. Covering UltrAuFoil films with a thin carbon layer recreates an adhesive surface, equivalent to that of pure carbon films (Figure 3, support 10).

Resistance to clipping was tested by simple examination at low magnification (Supplementary Figure S5). All commercial films showed multiple broken areas, leading to loss of sections in many cases. However, films with smaller holes and/or hole density (i.e. smaller observable surface) are more resistant, with R2/2 arrays more resistant than R3.5/1 and S7/2 (Supplementary Figure S4A-C). Strengthening of the films with the addition of an extra 30 nm carbon layer significantly improves resistance to clipping for all types of tested grids. Nearly 100% recovery of the grid surface was achieved with R2/1 (Supplementary Figure S5D). However, large size hole arrays (S7/2) remain fragile with increased probability of breaks. Strengthening of gold films with 30 nm carbon only slightly improved resistance to clipping, which suggest that mechanical properties of the film are dominated by those of the gold.

## DISCUSSION

In this work, we have established a simple protocol to determine at a glance which regions of the ribbon of section are suitable for best tilt series acquisitions. In addition to maximize the surface of adhesion, we have introduced a second electrical charging step during section collection. We also compared different support grids and selected those optimizing adhesion and/or resistance to clipping.

Low magnification tomographic reconstruction (Figure 1C-C”) and atlas recording at high defocus (Figure 1D, D’) both yield a complete adhesion map of the sections, the latter being altogether exhaustive and fast. They both show that cryosections deposited on the support film consist of a mosaic of adhesive and non-adhesive domains. Even for the best support films tested here, there exist non adhesive “bubbles” (Figure 1D, D’), not suited to tomographic acquisitions of even high quality 2D image recording. We suspect that this could be due to (i) a non-perfect flatness of the sections during generation, which could lead to local hills and valleys precluding “melting” propagation; and/or to (ii) the presence of adhesive and non-adhesive patches on the carbon film. Carbon films are indeed heterogeneous supports (Chu and Li, 2006) (Kim et al., 2021), which would explain why different results are obtained with different carbon sources (Figure 3). Quantifoil and homemade carbon surfaces reproducibly lead to the best adhesion, which is obtained for all samples in this case. Homemade continuous carbon films performed better than commercial continuous carbon from Quantifoil for 2D imaging.

Whereas the nature of the surface plays a role in adhesivity, the lattice type plays a role in (i) sample stability under the beam, and (ii) resistance to clipping. As mentioned above fre- standing sections over large holes are poorly stable under the beam and cannot be used directly. Free standing sections over small (2 µm) holes are also slightly less stable under the beam than sections adhering to a continuous film, leading to less driftless 2D images. This could be due to the presence of nano tears in some regions, which could propagate within free-standing sections and be fixed on adhesive films. We did not test the influence of the presence of a continuous film for tilt series acquisition on R2/1 or R2/2 lattices. But, with a very high yield of acquisition of tilt series suitable for tomographic reconstruction on bare R2/1 lattices, its absence can be expected of little consequence. In addition, it could introduce noise. To resist clipping reproducibly, commercial support films need to be thickened to 50 nm. In addition, the lattice type plays a role: the higher surface/hole ratio of the support film, the more resistant to clipping it is. S7/2 lattices present the highest observation surface (57% of the total surface), but - even thickened to 50 nm - remain fragile. They also often present long-range undulations which complicates tilt series acquisition. However, covered by an ultrathin continuous carbon film, S7/2 lattices, due to their large and adhesive imageable areas, are the best choice for 2D imaging in side-entry microscopes for which no clipping is required. R2/2 and R2/1 Quantifoil grids reinforced with carbon turned out to provide stable tilt series recording on well adhering sections. R2/1 lattices perform as well as R2/2 and should therefore be preferred on account of their higher imageable surface (26% *versus* 19%). Examples of 2D images and a reconstructed cryo-tomogram are shown in Supplementary Figures S6A and S6B respectively.

CEMOVIS is an alternative to cryo-FIB milling especially adapted to tissues and even whole organs or organisms which can be vitrified over large scale (typically up to 200 µm deep) in the absence of any cryoprotection by high pressure freezing. Another advantage is the possibility to generate ultrathin samples, well below 100 nm if desired (down to 20-30 nm), and serial sections. It can also be used as structure screening tool, with rapid sampling of large areas. Typically, a successful sectioning day results of 4 to 5 grids covered with a ∼ 2 *mm*-long and ∼ 50 *µm*-wide ribbon. Considering that 30% of the surface is lying over grid bars, about 70 to 90.10^3^ and 160 to 200.10^3^ *µm*^2^ are thus available for imaging on R2/1 and S7/2 grids respectively.

Together with stable generations of cryo-ultramicrotomes (Leica UC6/FC6 and UC7/FC7) and diamond knives (Diatome CEMOVIS 35 and 25), a series of progresses have been implemented in the CEMOVIS workflow over the years: (i) the use of double micromanipulators that although not fully automating the delicate generation and collection of ribbons of cryosections considerably alleviate the procedure (Hsieh et al., 2006)(Studer et al., 2014); (ii) the realization of the process inside a controlled environment; (iii) the use of an electrostatic press instead of a mechanical one (Pierson et al., 2010). To limit contamination, dry glove boxes have been used (Pierson et al., 2010) but introduce an extra obstacle in the handling of the sections. We opted for a partly dehydrated room, at 20 to 25 % relative humidity, which results in an optimal balance between contamination and comfort of the user. Cutting at the highest possible temperature below devitrification (-140°C) (Ng et al., 2020), use low angle knives (25°) (Han et al., 2008), and minimize the width of the block face (typically around 50-75 µm) (Al-amoudi et al., 2005) optimize section generation and minimize crevasses and/or compression. Our present work goes a step further, leading to high throughput cryo-ET. In our hands, the amount of tilt series suitable for optimal tomographic reconstruction was not different from that obtained with thin film samples. Typically, we obtained 50 tomograms per day, among which, more than 45 suitable for reconstruction.

This also opens a way toward serial tomography. We did not attempt here to record serial tomograms. In most samples, there are good chances that a given ROI found in an adhesive region of a section is also adhesive in the next section, as noticed by Ng et al. (2020). However, considering the available imaging surface of R2/1 lattices (26%), chances to access successive sections of a ROI are low. Highest observation surfaces are obtained using continuous support films, and carbon film-coated parallel-bar grids were successfully used by Ng *et al* to obtain serial tomograms of *S. cerevisiae* (Ng et al., 2020). However, these support films are likely to be too fragile to be clipped for tomographic acquisition in high end cryo-microscopes. S7/2 lattices tested here, even strengthened by additional carbon layer are still not resistant enough, and wavy, reducing high quality tomographic acquisition throughput. We did not test tilt series recording on sections deposited on Quantifoil R3.5/1 lattices (possibly covered with an additional ultrathin continuous film), but this could be an interesting solution, leading to higher probability of serial tomographic acquisitions.

## ACKNOWLEDGEMENTS

We thank Wim Hagen and Gurudatt Patra for help with data collection at the EMBL (Heidelberg, Germany) and the IGBMC (Illkirch, France) respectively. We were supported by the French Infrastructure for Integrated Structural Biology (FRISBI ANR-10-INSB-005) and Instruct-ERIC. FT and AL were supported by iNext-Discovery, grant number 871037, funded by the Horizon 2020 program of the European Commission.

