## Supplementary Figures for "Optimization of adhesion for high throughput cryo-electron tomography of vitreous sections"

### Supplementary Figure S1

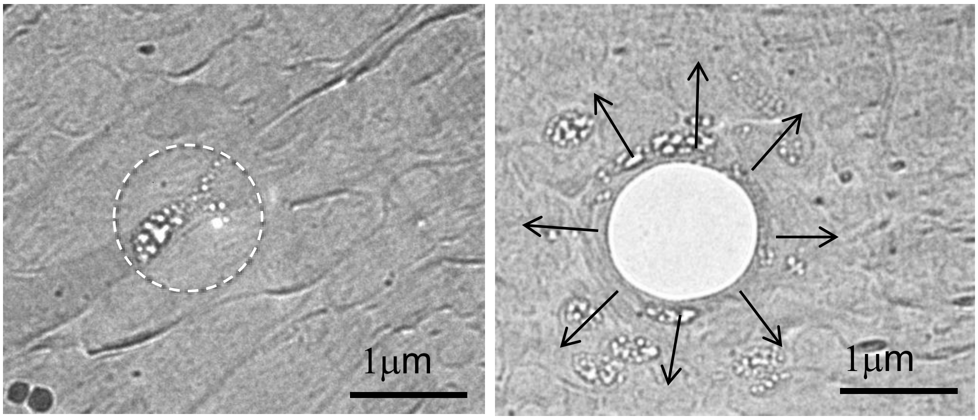


**Figure S1**. Focusing of the beam on an adhesive (left) and a non adhesive (right) region. Quantifoil S7/2 + 2 nm continuous carbon film.

### Supplementary Figure S2

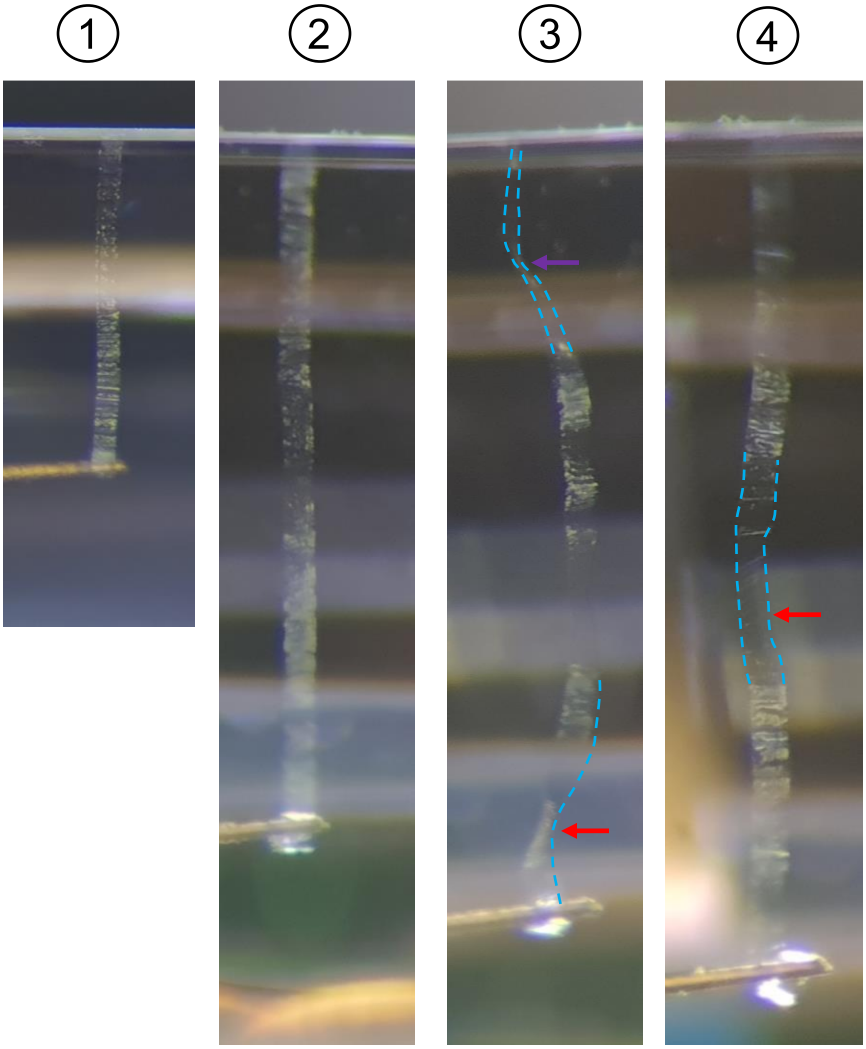


**Figure S2**. A few examples of ribbons during the generation process. Even and flat short ribbons are easily generated (1, 2). Longer ribbons (3, 4) tend to twist or bend.

### Supplementary Figure S3

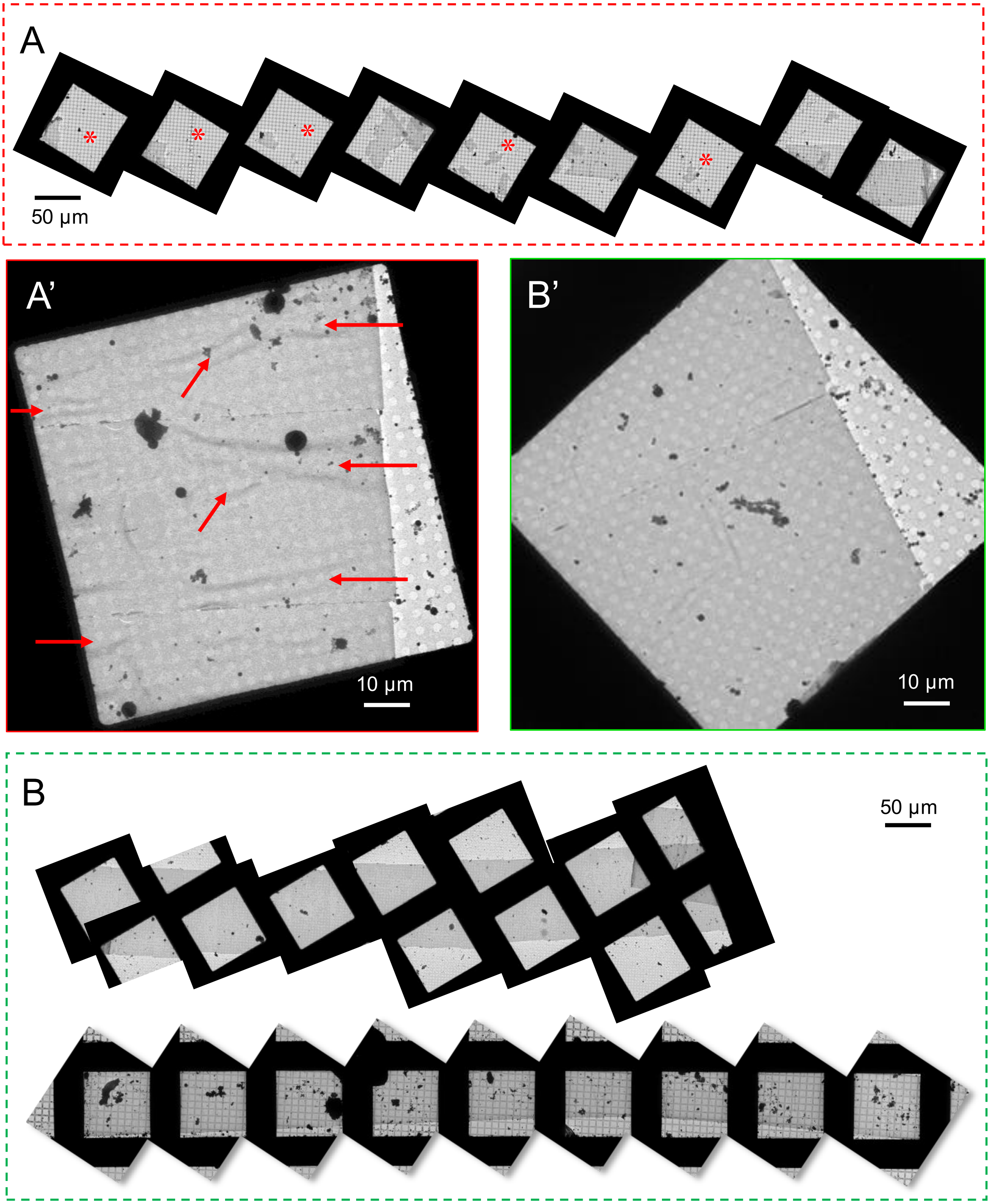


**Figure S3**. Cryo-EM low magnification imaging of ribbons collected with one-step (A, A') or two steps (B) electrical field charging (Crion electrode used in charging mode). In (A, A') missing regions resulting from large non-adhesive regions flown away by liquid nitrogen during transfer (red asterisks, A) and or numerous non-adhesive bubbles (red arrows, A') are observed. In (B, B'), ribbons are continuous and flat looking. All ribbons shown here were deposited on support films (Quantifoil R3.5/1 (A), R2/2 (A'B'), S7/2 (B)) covered by carbon evaporated from carbon threads in Leica ACE evaporator.

### Supplementary Figure S4

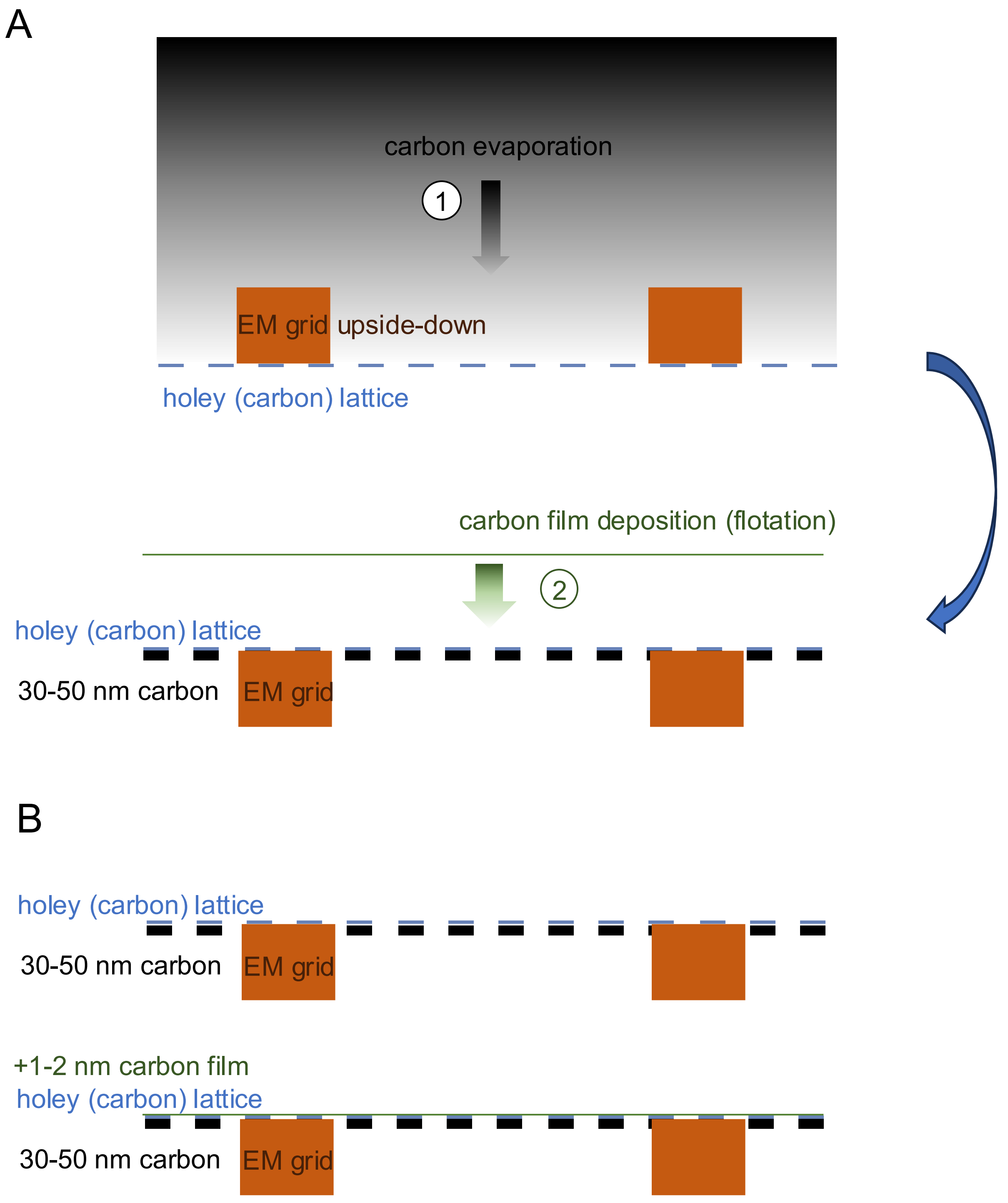


**Figure S4.** (A) Support film preparation, with strengthening of commercial support films by evaporation of an extra layer of carbon (1). On part of the grids, a continuous ultrathin carbon is then deposited by flotation (2). (B) Sketch of the two types of resulting support films.

#### Supplementary Figure S5

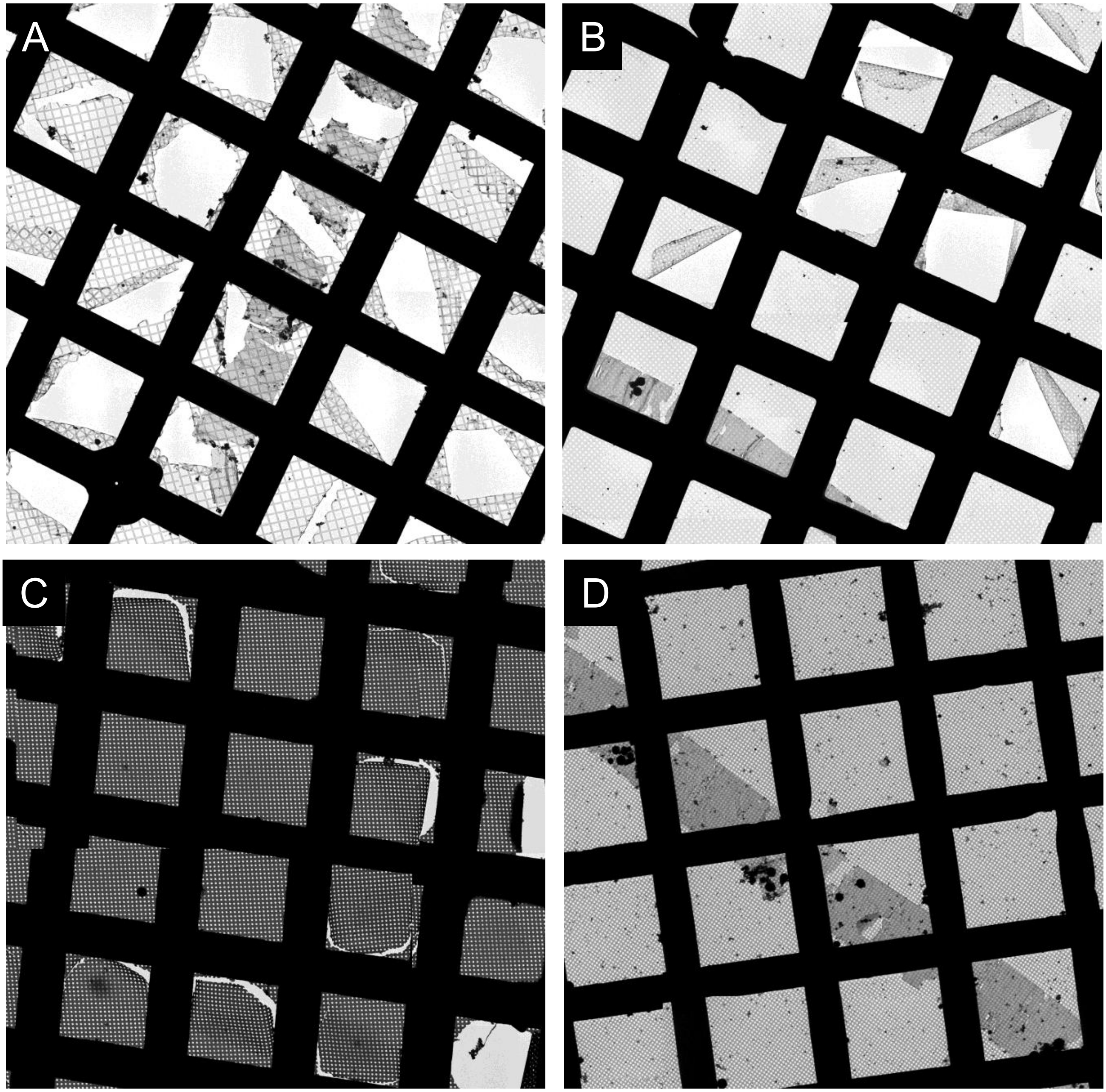


**Figure S5.** Examples of support grids after clipping, in the absence (A-C) or with (D) reinforcement with a 30 nm thick carbon layer. (A) Ribbon of cryosection deposited on Quantifoil S7/2 covered with an ultrathin carbon films (support 1 in Figure 3). (B) Ribbon of cryosection deposited on Quantifoil R2/2 (support 8 in Figure 3). (C) UltrAuFoil R2/2 (support 9). (D) Quantifoil R2/1 reinforced with a 30 nm carbon layer (support 6).

Supplementary Figure S6

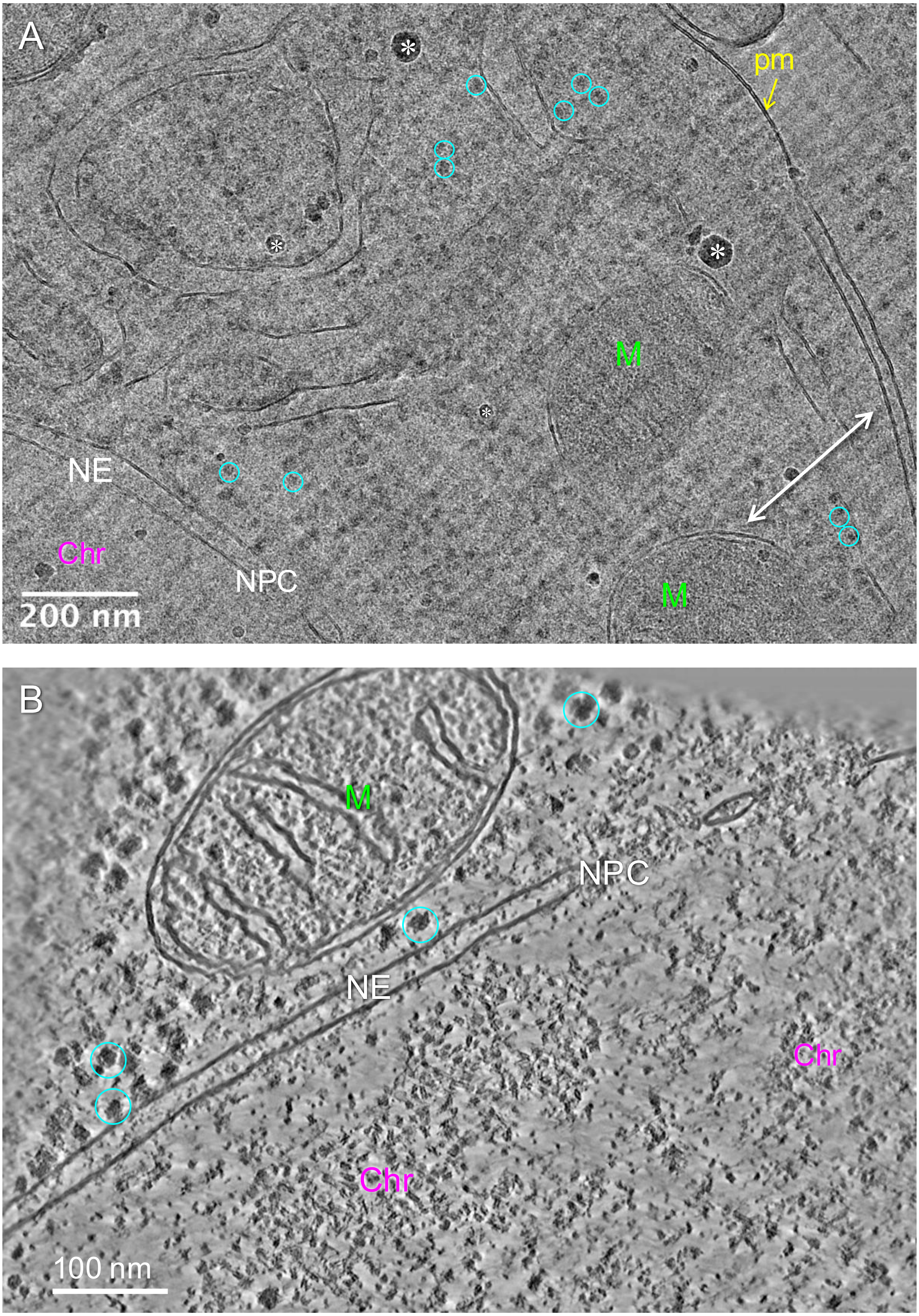


**Figure S6**. Examples of cryo-EM/ET imaging of the nuclear periphery of Drosophila embryonic brain. NE: nuclear envelope, NPC: nuclear pore complex, M: mitochondria, Chr: chromatin. A few ribosomes are circled in cyan. (A) Cryo-EM 2D image of a 50 nm-thick section. Some ice contamination deposited on the section surface are indicated by white asterisks. Cutting direction is indicated by the double white arrow. (B) Virtual slice (4.25 Å) in a cryo-tomogram of a 50 nm-thick section. Support grids: (A) Quantifoil S7/2 covered by 2nm-thick continuous carbon film; (B) Quantifoil R2/1 reinforced with 30 nm carbon.
